# Anthropogenic gradients shape *Staphylococcus/Mammaliicoccus* communities: Bacterial composition and resistance patterns as indicators of landscape hemeroby

**DOI:** 10.64898/2026.08.25.747043

**Authors:** Tamás Tari, Eszter Nagy, Olivér Lakat, Izabella Zám, Kimba Duncan Ombula, Brigitta Bóta, Rebeka Ráhel Nagy, Attila Zsolnai, Gábor Nagy, Ágnes Csivincsik

## Abstract

Antimicrobial resistance (AMR) is one of the greatest challenges within the One Health continuum. Exploring transmission routes between health domains and determining their driving forces are key priorities for future research. This effort can be effectively supported by landscape epidemiology, a field of science that integrates methods from landscape ecology and epidemiology to unravel the complex interdependencies behind disease transmission. This exploratory study aimed to demonstrate that landscape diversity and the degree of hemeroby (anthropogenic impact) correlate with the composition of bacterial communities and their AMR profiles. To test this hypothesis, submandibular lymph nodes from Cervidae and Suidae were collected to detect *Staphylococcus* and *Mammaliicoccus* bacteria and characterise their AMR features using selective culture and the VITEK 2 Compact automated system. As a result, the bacterial community in the more natural landscape was more diverse, characterised by the dominance of *Mammaliicoccus sciuri* and pan-susceptible isolates of *Staphylococcus hyicus*, and it displayed a low-level, heterogeneous AMR profile. Within the more hemerobic landscape, the bacterial community was characterised by the dominance of *Staphylococcus epidermidis*, a human-adapted species, and the AMR profile showed signs of higher antimicrobial pressure from both public health and veterinary origins. Although this study was based on only two study sites and was therefore not suitable for drawing definite conclusions, the findings suggest that human impact manifests itself in both bacterial and AMR profiles. A high prevalence of mammaliicocci and a heterogeneous AMR profile appeared to be indicators of naturalness. Conversely, the dominance of a human-adapted bacterial species and the accumulation of AMR features characteristic of medical environments likely indicate higher degrees of hemeroby.

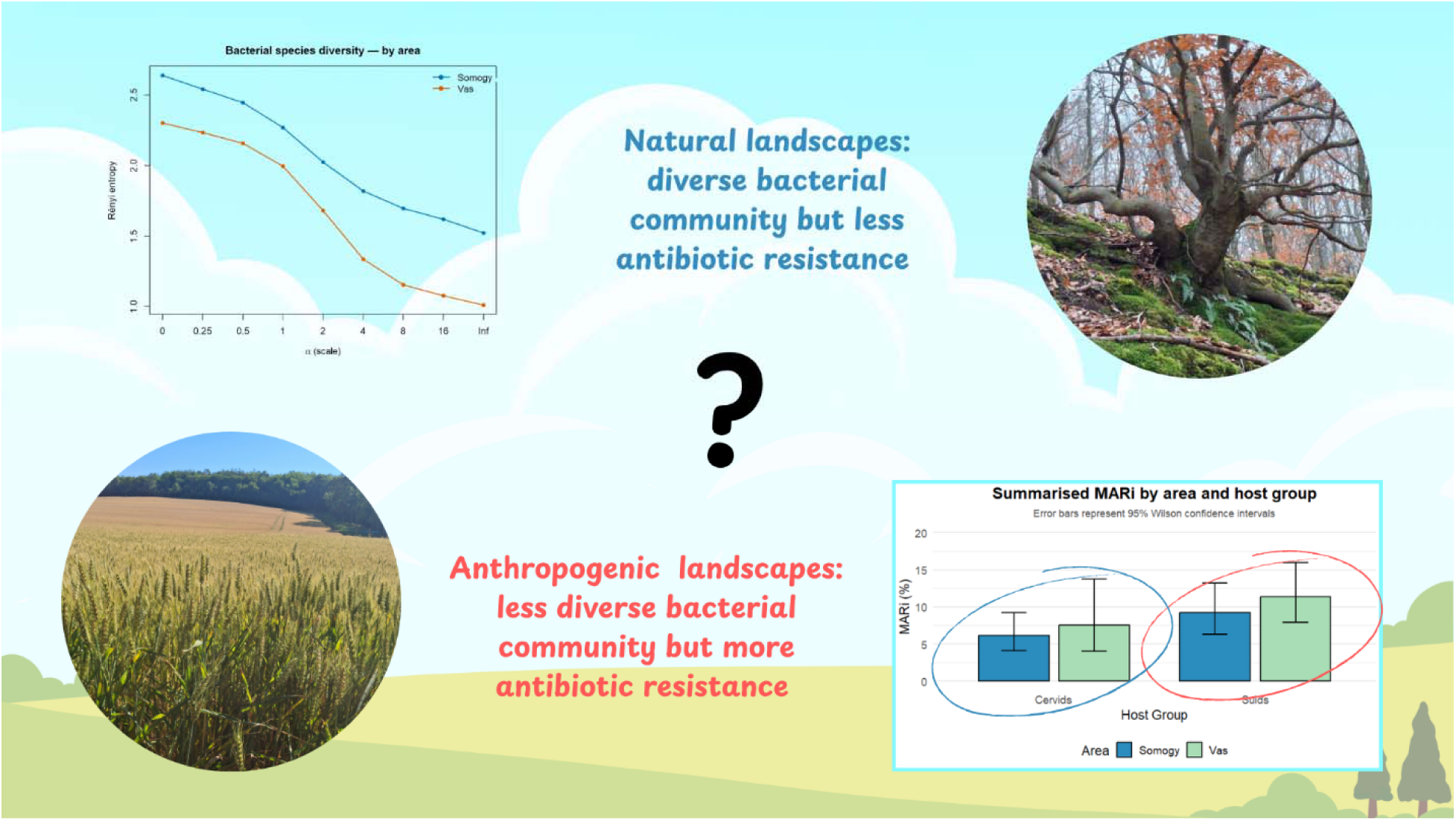

## Introduction

The central principle of One Health is that a well-functioning ecosystem guarantees the good health of humans, animals, and their environment (Adisasmito et al. 2022; Roy et al, 2025). Anthropogenic influence primarily erodes ecosystem functionality through the loss of naturalness and biological diversity (Ecke et al. 2025). Species richness ensures resilience against pathogens through the dilution effect, which means that in a diverse community, most species serve as dead-end hosts for pathogens, contributing to the fading out of epidemics (Da Costa et al. 2013). At the landscape level, hemeroby expresses the level of human impact (Sukopp, 1972; Walz & Stein 2014), while the diversity of land cover categories and the pattern of structural components contribute to the complexity of landscapes; therefore, in a multidisciplinary One Health study, evaluation of landscapes would be a part of the analysis (Hassel et al. 2021).

Antimicrobial resistance (AMR), though it is not a specific disease, is one of the main foci of One Health. Resistant bacteria and their genetic material circulate unimpeded across the human, animal, and environmental health domains (Medina et al. 2020). As a consequence, a growing proportion of the resistome (collection of resistance genes) in natural soil is of anthropogenic origin (Zhao et al. 2025). The frequent use of antimicrobials in human and veterinary medicine exerts artificial selection pressure on bacterial communities. Consequently, the frequencies of resistant genotypes increase, while susceptible ones die out (Mukherjee et al. 2021). Besides antimicrobials, other anthropogenic pollutants, such as heavy metals, pesticides, and other xenobiotics, can facilitate the drift toward an AMR crisis through co-selection. This phenomenon means that specific bacteria use the same mechanism to protect themselves from different xenobiotics; therefore, the evolution of resistance to any of them results in a general resistance to all of them (Despotovic et al. 2023). According to predictions, the current trends of AMR lead to a global health crisis by 2050. Both human and veterinary medicine will face diseases caused by multidrug-resistant bacteria, and consequently, an increasing number of fatal disease outcomes (Medina et al. 2020; Despotovic et al. 2023).

However, we know less about the environmental component of this process. A growing number of studies investigate AMR in wildlife (Vittecoq et al. 2016; Hassell et al. 2021; Plaza-Rodríguez et al. 2021; Ramos et al. 2022), but fewer of them analyse the potential resilience of certain habitats to the accumulation of AMR. Research findings indicate that higher levels of naturalness (Klümper et al. 2024) and landscape diversity (Da Costa et al. 2013; Sárközy et al. 2026) are associated with fewer AMR phenotypes in local bacterial communities. We presumed that a continental broad-leaved forest ecosystem was relatively resilient to AMR accumulation due to the biological diversity of its communities (Thonicke et al. 2020). Therefore, wild ungulates harvested in an area dominated by sylvatic habitats harbour fewer AMR phenotypes than those harvested in dominantly agricultural areas. To test our hypothesis, we collected submandibular lymph nodes from legally hunted, visibly healthy ungulate carcasses. These lymph nodes drain the area of the oral and nasal cavities and the facial region, filtering the pathogens invading the inner tissues (von Bargen et al. 2015). The presence of cultivable bacteria here, even in the absence of any visible lesions, is well documented (Hanlon et al. 2016; Ravel et al. 2015).

The members of the genera *Staphylococcus* and *Mammaliicoccus* are good model bacteria for AMR research. These microbes are ubiquitous, frequently occurring on the skin (Wesołowska & Szczuka 2023), in the intestinal tract (Rahman et al. 2026), the nasal cavity (Schauer et al. 2021), and in the environment of animals (Martins-Silva et al. 2023). In addition, the selective culture of these bacteria is based on their halophilic features. Supplementing culture media with up to 15% NaCl successfully impedes the growth of concurrent bacteria, enabling investigators to obtain pure cultures almost in one step (Kročko et al. 2019; Onyango & Alreshidi 2018). Automated systems with reliable accuracy, such as VITEK Compact 2 of BioMérieux, are available for the identification and antimicrobial susceptibility testing of many members of the *Staphylococcus* and *Mammaliicoccus* genera (Bobenchik et al. 2014).

For comparing the diversity of landscapes, bacterial communities, and AMR phenotypes between study areas, the Rényi diversity profile analysis offers a good opportunity. In contrast with most single-value diversity indices, this method provides a comprehensive insight into the structural complexity of the investigated biological system (Tóthmérész 1993; Ricotta et al. 2021). Besides diversity, landscape patterns (Hesselbarth et al. 2019) and hemeroby (Walz & Stein, 2014) can contribute to the resilience of a specific landscape; therefore, the calculation of landscape metrics would support the epidemiological analysis.

By selective culture of *Staphylococcus* spp. and *Mammaliicoccus* spp. bacteria from the submandibular lymph nodes of visibly healthy animals from two different study areas that were compared by Rényi diversity profiles, landscape patterns, and hemeroby, this study aimed to answer the following research questions:

● Is supplementing culture broth with 10% NaCl suitable to obtain bacteria of the *Staphylococcus* and *Mammaliicoccus* genera from tissue samples?
● Is the prevalence of *Staphylococcus/Mammaliicoccus* spp. in the submandibular lymph nodes of healthy ungulates (wild boar, red deer, fallow deer) high enough to use it for further statistical analysis?
● How diverse are the *Staphylococcus/Mammaliicoccus* communities of the study areas?
● How diverse are the AMR phenotype compositions of the study areas?
● Is there any difference between the host species relating to bacterial and AMR diversity?
● Is there any correlation between the effects of hosts and study areas on either bacterial diversity or AMR profile?
● Do bacterial diversity or AMR profiles reflect landscape diversity profiles, landscape structure, and hemeroby?

## Materials and methods

### Study areas

The two study areas are located in Transdanubia in Hungary, adjoining eastern Croatia and the Pannonian Region of Slovenia along the River Drava and River Mura, respectively. The three countries share a special biogeographical region with a diverse landscape mosaic featuring flat plains, rolling hills, and low mountains (Reed et al. 2004). The sub-Mediterranean and sub-Atlantic climates affect this area, resulting in a transition zone where Western Balkan (Illyrian) plant species coexist with Pannonian flora (Salamon-Albert et al. 2011; Fekete et al. 2014). This part of Hungary is one of the most forested areas supporting dense populations of wild ungulates and the golden jackal (*Canis aureus*) (Kemenszky et al. 2022). Within the study area, a remarkable zonation can be observed in both climate and vegetation from southwest to northeast due to the dual climatic influence.

The Somogy study site (9398 ha) is located in Somogy County, in a hilly area with a cooler and more humid local climate, which results in moisture-indicating forest communities, such as oak-hornbeam mixed forests and beech stands, particularly in deep valleys and on the northern slopes. The largest silver lime (*Tilia tomentosa*) population of Hungary lives here. Most of the study area is covered by forests. Contrary to the humid climate, permanent surface waters are rare within the territory. Wildlife management is characterised by big game, such as wild boar (*Sus scrofa*), red deer (*Cervus elaphus*), roe deer (*Capreolus capreolus*), fallow deer (*Dama dama*), and mouflon (*Ovis gmelini*). The majority of the site comprises the Zselic Landscape Protection area with an Important Bird Area (IBA). Agricultural activity is limited to grassland and cropland management and sheep farming on the edge of the territory. The Vas study site (3596 ha) is in Vas County, in the northwestern part of the Transdanubian hilly area; therefore, it is under milder Mediterranean and stronger Atlantic climate influence, compared to the Zselic area. Although this area is also dominated by woodland habitats, the proportion of agricultural landscapes is relevant, primarily in the northern part of the area. This territory is rich in surface waters. Wildlife management is based on dense wild boar and red deer populations. The Vas site does not contain a protected area. Agricultural activity is characterised by cropland cultivation and cattle grazing (Figure 1).

**Figure 1.**
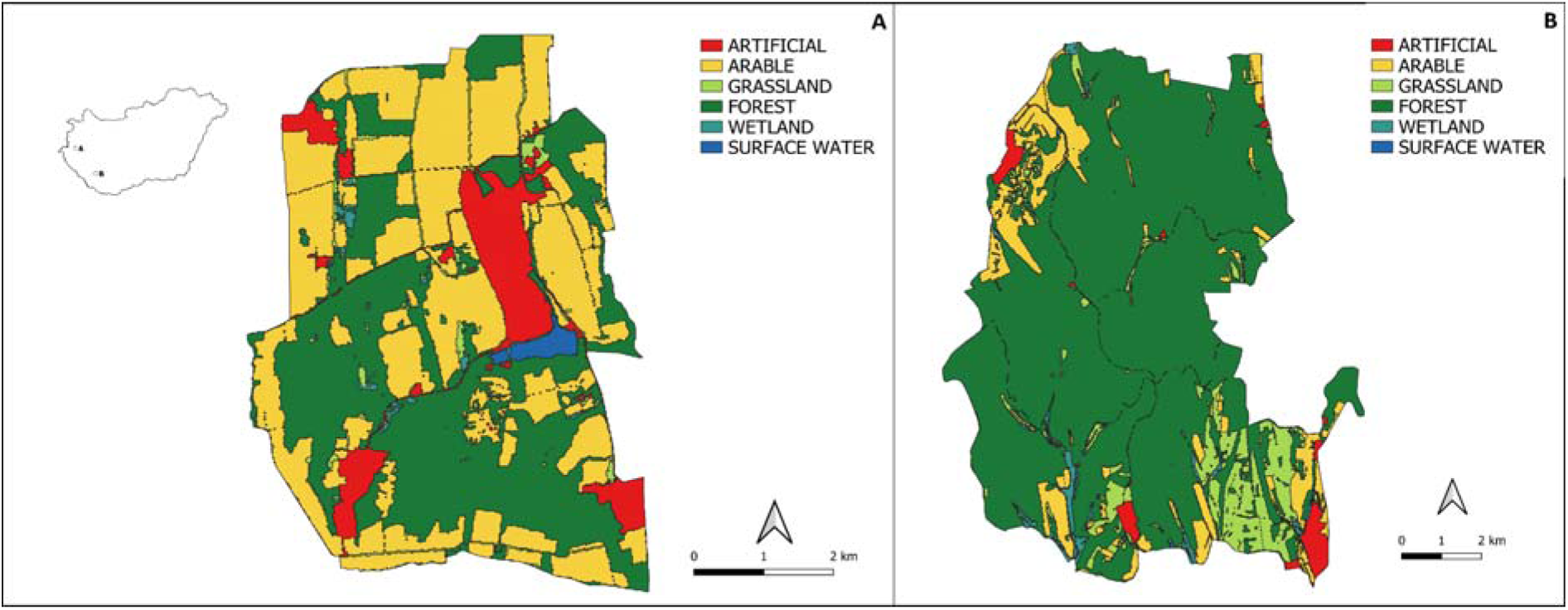
Maps of the two study sites (A: Vas, 3596 ha; B: Somogy, 9398 ha) and their location in Hungary (small map). Figure legend: ARTIFICIAL - built-up surfaces, ARABLE - arable land, GRASSLAND - meadows and pastures, FOREST - native woodlands and non-native tree plantations, WETLANDS - wet grasslands, moors, and alder carrs, SURFACE WATER - brooks and ponds.

### Sample collection

The carcasses in this study originated from legal hunting activities and were not harvested for scientific purposes. In Hungary, the Act LV of 1996 on “Wildlife protection, wildlife management, and hunting” (see https://njt.jog.gov.hu/jogszabaly/1996-55-00-00, in Hungarian, accessed on 06 July 2026) determines the rules on wildlife management. Section 47. § of this Act imposes an obligation on the game manager of the hunting territory to prepare an “annual wildlife management plan” and submit it to the hunting authority for approval. All sampled carcasses were taken within the framework of the officially approved annual wildlife management plans of the respective hunting territories.

The sample collection included wild boar (Suidae), red deer and fallow deer (Cervidae) carcasses. Following routine evisceration of the carcasses, the submandibular lymph nodes were removed along with a 2-3 mm thick layer of connective tissue to avoid contaminating the organs. The collected lymph nodes were placed into individually marked disposable plastic bags and stored in a cooler for transportation to the laboratory, where the specimens were immediately deep-frozen.

### Laboratory work

The collected lymph node specimens were stored at -20 °C until processing. After an approximately 60 min thawing period, the surface of the specimen was flamed, and it was put into a sterile porcelain mortar. Using a sterile blade to cut and sterile sand to grind, we produced a tissue mince, which was then mixed with 10 mL of buffered peptone water liquid medium supplemented with 10% sodium chloride (BPW-10%). This medium was produced in-house from Buffered Peptone Water, GranuCult® Prime; Sigma-Aldrich, Merck KGaA, Darmstadt, Germany. The inoculated broth media were incubated at 37 °C for 24 hours to ensure bacterial revival. From each tube of BPW-10%, 100 μL inoculum was spread on the surface of Plate Count Agar (PCA; prepared in-house with Plate Count Agar, GranuCult® Prime; Sigma-Aldrich, Merck KGaA, Darmstadt, Germany, in accordance with the manufacturer’s instructions). Inoculated PCA plates were incubated for 48 hours at 37 °C and were checked every 24 hours.

We selected suspect *Staphylococcus* colonies for transfer to the next PCA medium for purification. A colony was chosen if it was 1-3 mm, round, convex, and opaque, with a smooth surface and entire edge (Shaw 1951; Agarwal et al. 2022). Pure cultures were obtained using a four-quadrant streak plate method. Gram staining was performed on pure cultures using the Gram-Nicolle Kit (VWR International, LLC, Radnor, PA, USA) in accordance with the manufacturer’s instructions to differentiate between Gram-positive and Gram-negative bacteria.

All bacterial isolates, which were Gram-positive cocci, were processed using the VITEK Compact 2 system (bioMérieux, Marcy-l’Étoile, France) with VITEK 2 GP (Ref. No. 21342) Gram-positive identification cards. Antimicrobial susceptibility testing (AST) was carried out on bacterial isolates identified with at least 90% probability. For AST, we used VITEK 2 AST-P592 (Ref. No. 222887) card, which is suitable to test antimicrobials as follows: cefoxitin (FOX), benzylpenicillin (B-PEN), oxacillin (OXA), gentamicin (GEN), ciprofloxacin (CIP), moxifloxacin (MOX), erythromycin (ERY), clindamycin (CLIN), linezolid (LIN), teicoplanin (TEI), vancomycin (VAN), tetracycline (TET), tigecycline (TIGE), fosfomycin (FOM), fusidic acid (FUS), rifampicin (RIF), and trimethoprim (TRIM). Findings were exported to Excel sheets (Microsoft® Excel® for Microsoft 365 MSO, Version 2511).

External reference strains were not processed in parallel with test specimens; therefore, strict internal validation criteria were applied. Isolates of a “Good” or higher (“Very good” or “Excellent”) confidence level in identification were included, while others were excluded, ensuring the validity of the AMR data. Furthermore, all inocula were standardised to a turbidity of 0.5–0.63 McFarland via the VITEK DensiCHEK device (bioMérieux, Marcy-l’Étoile, France), and antibiograms validated by the VITEK 2 Advanced Expert System (AES) were reported as ‘Consistent’, ‘Consistent with correction’ and ‘Inconsistent’. These remarks were added to AMR results.

Notably, this automated system was developed for clinical use; therefore, it is optimised for pathogenic bacteria of human medicine. For this reason, its database contains the biochemical features and antimicrobial phenotypes of few environmental bacteria. Furthermore, its diagnostic accuracy is lower than that of the gold standard methods (Bobenchik et al. 2014; Adade et al. 2024).

### Data processing

Laboratory findings were exported to Excel sheets (Microsoft® Excel® for Microsoft 365 MSO, Version 2511). All analyses were performed in an R 4.6.0 environment, using readxl (Wickham and Bryan 2025), vegan (Oksanen et al. 2026), ggplot2 (Wickham 2016) and tidyr (Wickham et al. 2025) packages. Permutation tests were run with 9999 repetitions. The study was exploratory and hypothesis-generating; accordingly, it was not adjusted for multiple testing, and the individual comparisons are interpreted based on the raw p-values in an exploratory manner (Rothman 1990; Bender and Lange 2001).

The distribution of bacterial species by area and by game taxa was analysed using contingency table methods. The game taxon and the area were crossed (both game taxa occurred in both areas), so the area and game taxon effects did not coincide, and the two factors were evaluated separately. For the species × area and the species × wild taxon tables, a chi-squared test with Monte Carlo-based p-value (9999 repetitions) was applied, and the 2×2 comparisons per species were made with a Fisher’s exact test.

Resistance profiles were treated as binary (resistant/sensitive) vectors per isolate. The 8 antibiotics to which no isolate was resistant (invariant, zero-variance variables) were excluded from the analysis, and the remaining 9 variable antibiotics were used. Differences between isolates in pairs were expressed as Jaccard distance; isolates that did not show resistance (pan-sensitive) were considered mutually identical (distance = 0). The resistance profiles were treated as binary (resistant/sensitive) vectors per isolate. Differences between isolates in pairs were expressed as Jaccard distance; isolates that did not show resistance (pan-susceptible) were considered mutually identical (distance = 0).

The pattern of antibiotic resistance by species was displayed in a heatmap, where colour indicates the ratio of resistant isolates (0–1), and cell values show the number of resistant isolates per species and antibiotic. Data from all 17 antibiotics tested were used for the representation.

The composition of the resistance profile was investigated by a two-way permutation multivariate analysis (PERMANOVA; vegan::adonis2) (Anderson 2001; Oksanen et al. 2026), including the area (Somogy/Vas) and the wild taxon (Suidae/Cervidae) and their interaction as explanatory variables. The homogeneity of the multivariate dispersion between the groups was verified by the PERMDISP test (vegan::betadisper, permutest) with the same (Jaccard) distance measure. The profile patterns were displayed by principal coordinate analysis (PCoA; cmdscale), coloured by area. Differences in individual antibiotic levels between the areas were examined for the 9 variable antibiotics using Fisher’s exact test.

The prevalence of *Staphylococcus/Mammaliicoccus* infection was calculated as the proportion of samples that produced an identifiable *Staphylococcus/Mammaliicoccus* isolate. If any of the samples had more than one isolate of the same bacterial species, only one was analysed further to avoid pseudoreplication. Prevalence of AMR was calculated as the proportion of isolates that were resistant to at least one tested antimicrobial. Prevalence of multi-drug resistance (MDR) was determined as the proportion of isolates that were resistant to at least three drug classes (e.g., beta-lactams, macrolides, tetracyclines…). The Multiple Antimicrobial Resistance index (MARi) was calculated as the proportion of positives among all completed tests. This index was determined for specific isolates as the number of AMR phenotypes divided by the number of tests in the card (N=17). For the study areas and the host groups, MARi was calculated as the total number of AMR phenotypes detected in the specific study unit divided by the total number of completed AST tests (17 × the number of isolates). Prevalence of *Staphylococcus/Mammaliicoccus* infection, AMR, MDR, and MARi were compared between study areas and host groups using Fisher’s exact test. The results of these analyses were visualised using the ggplot2 (Wickham 2016) package.

The land cover of sampling areas (hunting areas), bacterial species diversity, and antibiotic resistance were characterised by Rényi’s diversity profiles. In this method, diversity is plotted on the entire α scale (α = 0–∞), where α = 0 corresponds to the type richness, α = 1 corresponds to the Shannon index, α = 2 corresponds to the inverse Simpson index, and α → ∞ reflects the weight of the dominant type (Hill 1973; Tóthmérész 1995), as follows:

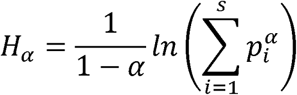

where Hα is the Rényi entropy, α is the scale parameter, and pi is the relative frequency of a given land cover type or the number of isolates belonging to a specific bacterial species or AMR phenotype. The profiles were prepared by area (Somogy/Vas) and by the host group (Suidae/Cervidae). If the profile of one group is above that of another across the entire scale, it is clearly considered more diverse; if the profiles intersect, the diversity ranking is considered α-dependent.

The landscape structure of the two study areas was characterised using the 20 m-resolution, three-level nomenclature coverage grid of the Ecosystem Base Map (http://alapterkep.termeszetem.hu/, accessed on 6 December 2025). The analysis was carried out at two thematic breakdowns: (i) at level 1 of the map, aggregated into the six main categories (artificial surfaces, croplands, grasslands and other herbaceous plants, forests and other woody plants, wetlands, surface waters), and (ii) at level 3, at the level of detailed habitat subtypes, particularly with a focus on forest typology. The compositional diversity of the landscapes was characterised by the Rényi diversity profile calculated from the territorial proportions of the coverage categories (Hill 1973; Tóthmérész 1995), on both thematic levels. The spatial arrangement (configuration) of the landscape structure was expressed by patch density (PD; patches/100 ha) and edge density (ED; m/ha) at both levels. The patches were demarcated based on eight neighbourhoods; the landscape boundary was not counted as an edge. The calculations were performed using the terra (Hijmans 2025) and landscapemetrics (Hesselbarth et al. 2019) R packages.

The naturalness of the areas was characterised following the hemeroby concept for landscapes (Sukopp 1972). The subtype-level coverage classes were classified into naturalness degrees, adapting hemeroby classification by Walz and Stein (2014), as follows: 1 = metahemorobic and polyhemorobic, 2 = α-euhemorobic, 3 = β-euhemorobic A, 4 = β-euhemorobic B, 5 = oligohemorobic (Table 1).

**Table 1.** Classification of naturalness adapted from the work of Waltz and Stein (2014).

| n (level of naturalness) | Land cover categories | Reason for classification |
| --- | --- | --- |
| 1 (meta/polyhemorobic) | Artificial surfaces | Built-up and paved area - very strong human impact |
| 2 (α-euhemorobic) | Agricultural lands, non-native woodlands | Extensively managed - strong human impact |
| 3 (β-euhemorobic A) | Grasslands and shrublands | Semi-natural with moderately strong human impact (grazing) |
| 4 (β-euhemorobic B) | Surface waters | Semi-natural with moderate human impact (flood control, aquaculture) |
| 5 (oligohemorobic) | Native forest habitats, wetlands | Semi-natural with weak human impact (forests available for wood supply) |

The average naturalness of the study areas was calculated using the equation

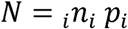

where N is the average naturalness, n is the degree of naturalness of the i landscape category, p is the proportion of the i landscape category within the study site.

Contrary to the relatively high biodiversity of broad-leaved forests, particularly within the landscape protection area at the Somogy site, the ahemorobic category by Walz and Stein (2014) was not applied, since the forests in the studied regions of Hungary are classified into archetypes from medium to very high-intensity use (Barredo et al. 2025). Because there are only two areas, the comparison of landscape structure and naturalness was interpreted descriptively and in a hypothesis-generating manner, without a statistical test at the landscape level.

## Results

This study investigated 82 lymph node specimens from two different study areas and two groups of host species. From a Somogy County hunting estate (Somogy), 30 Cervidae (19 red deer, 11 fallow deer), and 22 Suidae (wild boar) samples were collected, while from the Vas County area (Vas), 15 red deer and 15 wild boar samples were collected. Following laboratory processing, a total of 77 *Staphylococcus* (N = 63) and *Mammaliicoccus* (N = 14) isolates were obtained, of which 57 were suitable for antimicrobial susceptibility testing. All AST-tested bacterial isolates were successfully tested for all 17 antimicrobials on the AST card used, resulting in 969 AMR phenotype characteristics. The complete dataset of the study is available at https://zenodo.org/records/22096010.

Visual comparison of the *Staphylococcus/Mammaliicoccus* community in the two areas, Somogy appears to have higher diversity of bacteria than the Vas area (Figure 2. A). Of the two host groups, Suidae showed a higher potential in accumulating a diverse bacterial community (Figure 2. B).

**Figure 2.**
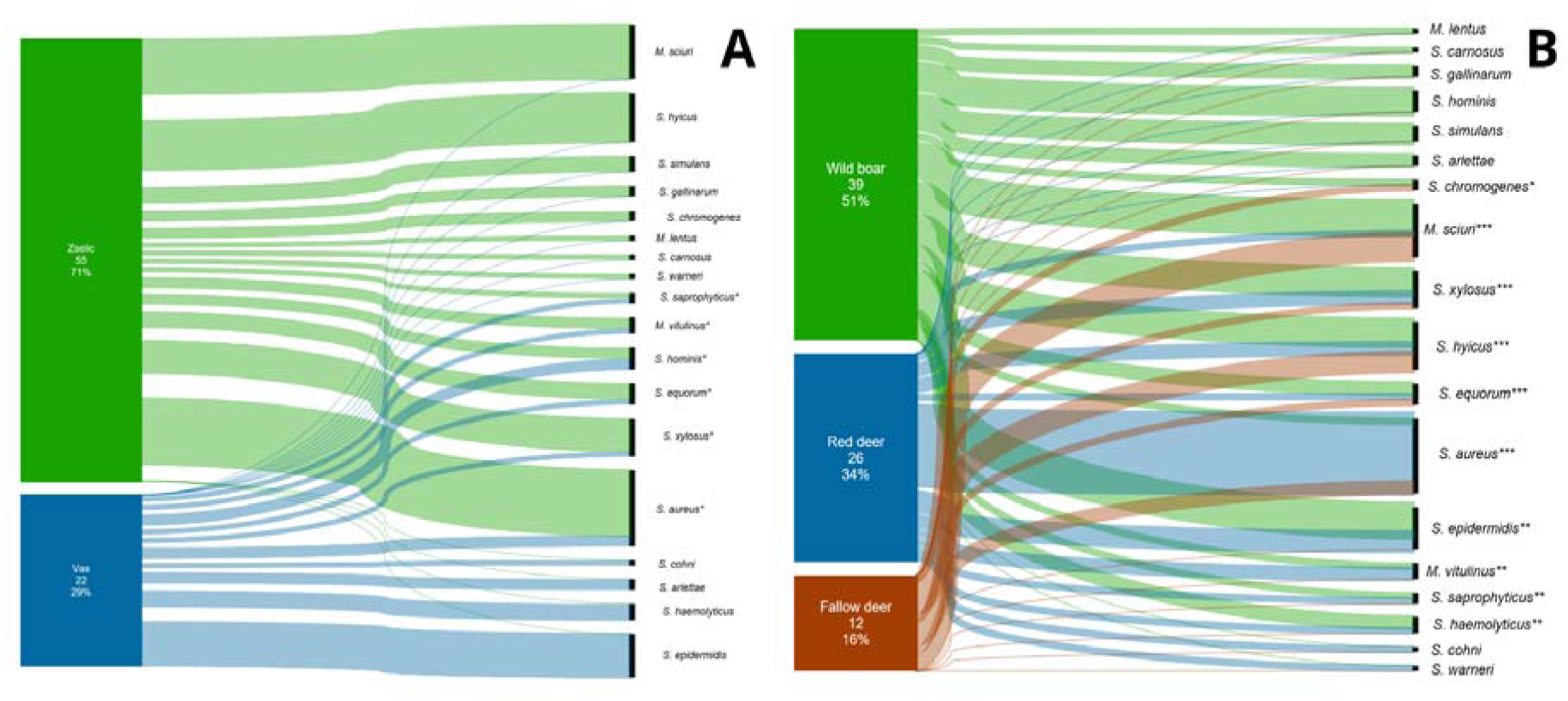
The composition of the *Staphylococcus/Mammaliicoccus* community at the two study sites (A) and in the host species (B) visualised using Sankey diagrams.

*Mammaliicoccus* isolates came predominantly from Somogy (13/14), while only one *M. vitulinus* isolate was found in Vas. The most common *Mammaliicoccus* species was *M. sciuri* (10 isolates), which occurred only in Somogy. Somogy isolates were apparently dominated by *S. aureus*, *M. sciuri*, and *S. hyicus*, while the predominant species of the Vas territory was *S. epidermidis*.

The species composition was significantly different between the two areas (chi-squared Monte Carlo test: χ² = 50.11, p = 0.0001). According to the species-specific Fisher’s exact test, the strongest, area-bound species was *S. epidermidis*, which occurred exclusively in Vas (0 Somogy / 8 Vas; p < 0.001); as a weaker sign, *S. haemolyticus* was also Vas-specific (0 / 3; p = 0.021), while the attachment of *S. hyicus* (9 Somogy / 0 Vas; p = 0.053) and *M. sciuri* (10 / 0; p = 0.054) to Somogy remained borderline. The distribution of the other species did not differ substantially (Table 2).

**Table 2.** Bacterial species isolated at the two study sites, Somogy and Vas, from the submandibular lymph node specimens of wild ungulates.

| Bacterial species | Number of isolates in Somogy | Number of isolates in Vas | Total number of isolates |
| --- | --- | --- | --- |
| *Mammaliicoccus lentus* | 1 | 0 | 1 |
| *Mammaliicoccus sciuri* | 10 | 0 | 10 |
| *Mammaliicoccus vitulinus* | 2 | 1 | 3 |
| *Staphylococcus arlettae* | 0 | 2 | 2 |
| *Staphylococcus aureus* | 12 | 2 | 14 |
| *Staphylococcus carnosus* ssp.<br>*carnosus* | 1 | 0 | 1 |
| *Staphylococcus chromogenes* | 2 | 0 | 2 |
| *Staphylococcus cohnii* ssp.<br>*urealyticum* | 0 | 1 | 1 |
| *Staphylococcus epidermidis* | 0 | 8 | 8 |
| *Staphylococcus equorum* | 3 | 1 | 4 |
| *Staphylococcus gallinarum* | 2 | 0 | 2 |
| *Staphylococcus haemolyticus* | 0 | 3 | 3 |
| *Staphylococcus hominis* ssp. *hominis* | 2 | 2 | 4 |
| *Staphylococcus hyicus* | 9 | 0 | 9 |
| *Staphylococcus saprophyticus* | 1 | 1 | 2 |
| *Staphylococcus simulans* | 3 | 0 | 3 |
| *Staphylococcus warneri* | 1 | 0 | 1 |
| *Staphylococcus xylosus* | 6 | 1 | 7 |
| **Sum** | **55** | **22** | **77** |

The species composition also differed between game taxa (chi-squared Monte Carlo test: χ² = 26.70, p = 0.023), but a single species drove it: *S. aureus* showed a strong preference for Cervidae (13 Cervidae / 1 Suidae; p < 0.001). None of the other species differed significantly among the game taxa (Table 3). The game taxon was crossed with the area (Suidae and Cervidae both occurred in both areas: Somogy 31/24, Vas 7/15), so the area and game taxon effects did not coincide.

**Table 3.**
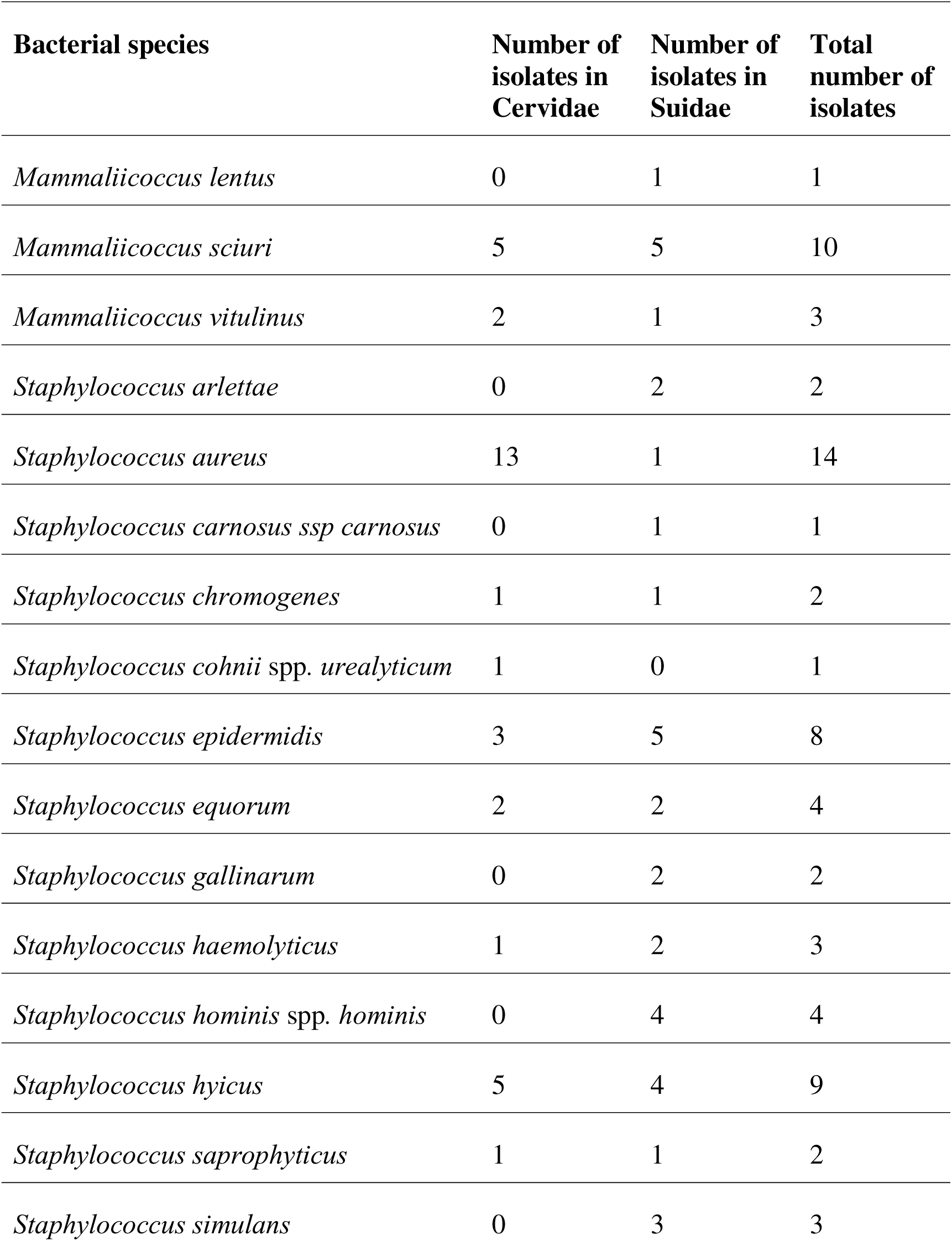
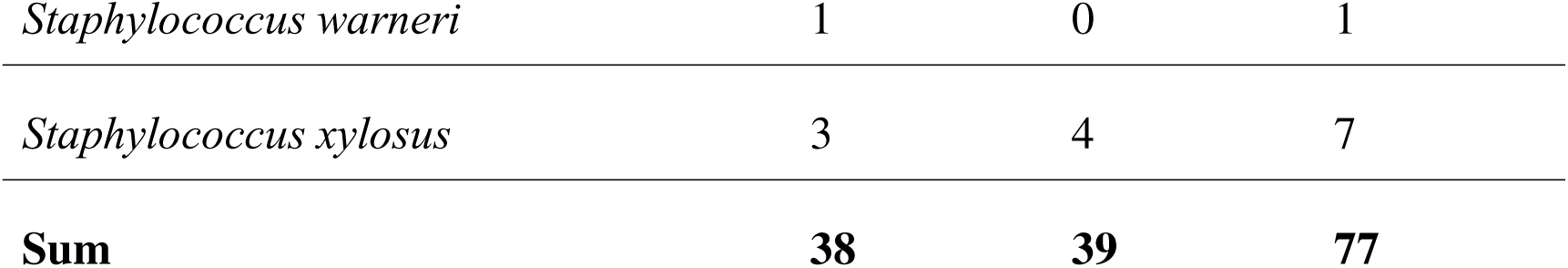
Bacterial species isolated from the submandibular lymph node specimens of the two host groups, Cervidae and Suidae.

The composition of the resistance profile differed significantly between the two hunting areas (two-way PERMANOVA: F = 8.28, p = 0.0002, R² = 0.13), but not according to the game taxon (F = 0.71, p = 0.52) and the area × game taxon interaction (F = 0.59, p = 0.62). The multivariate dispersion differed significantly between the areas over the entire dataset (PERMDISP: F = 5.73, p = 0.021); Somogy was more heterogeneous (average centroid distance 0.54 vs 0.46). This result confirmed that the difference in the resistance profile between the areas was due to real compositional (location) differences, not to the heterogeneity of the scatters (Table 4).

**Table 4.** Antimicrobial resistance (AMR) phenotypes occurred at the two study sites, Somogy and Vas. The numbers and the proportion of resistant isolates among all antimicrobial susceptibility-tested isolates are presented. An asterisk marks the significant difference in the frequency of a specific AMR phenotype between the two sites.

| Antimicrobial | Resistant isolates in Somogy | Resistance % in Somogy | Resistant isolates in Vas | Resistance % in Vas | p-value |
| --- | --- | --- | --- | --- | --- |
| B-PEN | 14 | 39 | 17 | 81 | 0.00261* |
| TET | 0 | 0 | 5 | 24 | 0.00486* |
| FUS | 16 | 44 | 3 | 14 | 0.02295* |
| ERY | 0 | 0 | 2 | 10 | 0.13158 |
| FOM | 4 | 11 | 4 | 19 | 0.44903 |
| CLIN | 4 | 11 | 4 | 19 | 0.44903 |
| CEF | 2 | 6 | 1 | 5 | 1.0 |
| OXA | 2 | 6 | 1 | 5 | 1.0 |
| MOX | 1 | 3 | 0 | 0 | 1.0 |

The main coordinate analysis of the resistance profiles (PCoA, Jaccard distance) illustrated the partial, overlapping separation of the areas with centroid shifts, consistent with the PERMANOVA result (Figure 3); significant overlap between the groups was consistent with the low explained variance (R² = 0.13).

**Figure 3.**
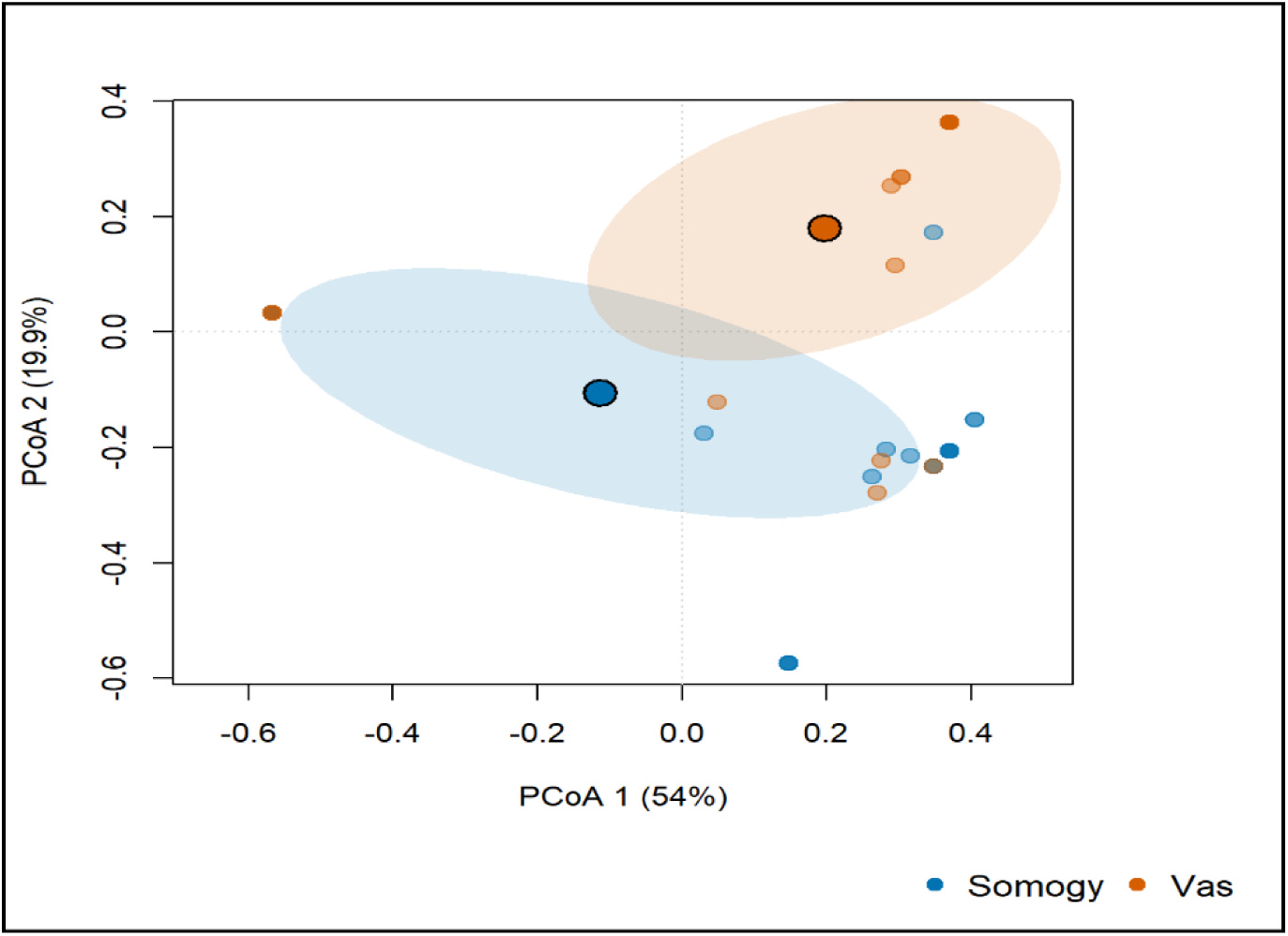
Principal coordinate analysis (PCoA) of antibiotic resistance profiles of *Staphylococcus* and *Mammaliicoccus* isolates based on Jaccard distance across 9 antibiotic variables. The dots mark isolates, coloured by area (blue = Somogy, orange = Vas); the large circles with black borders show the regional centroids, and the shaded ellipses show the scattering of the groups (±1 SD). The percentages next to the axes indicate the variance explained. The overlapping ellipses reflect partial separation, consistent with the significant but moderate area difference (PERMANOVA).

According to the pairwise Fisher’s exact test, the regional difference was primarily driven by three antibiotics. Two showed dominance in Vas: B-PEN (Somogy 25% vs Vas 85%; p < 0.001) and TET (0% vs 25%; p = 0.009), while the third was characteristic of Somogy (FUS 44% vs 14%; p = 0.023) and was primarily associated with *M. sciuri* isolates.

The heatmap of AMR patterns illustrates that most of the investigated antimicrobials are effective against all isolated bacteria. The most frequent AMR phenotypes were B-PEN, FUS, FOM, and CLIN, while the bacterial species that possessed the most AMR phenotypes were *M. sciuri*, *S. epidermidis*, and *S. xylosus* (Figure 4).

**Figure 4.**
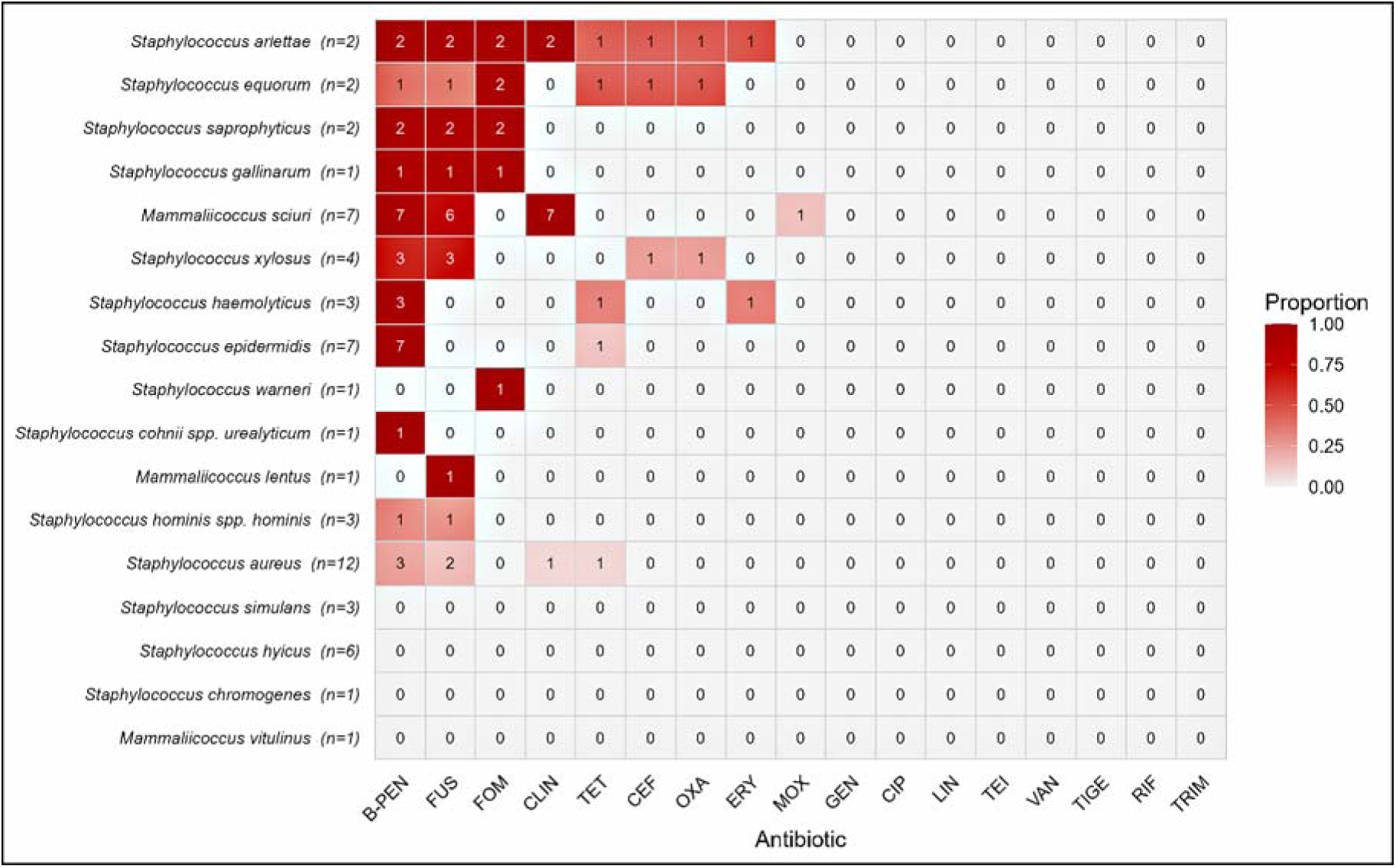
Heat map of antibiotic resistance in *Staphylococcus* and *Mammaliicoccus* species. Colour indicates the ratio of resistant isolates within a given species (0–1; light grey = 0, dark red = 1); the numbers in the cells are the counts of resistant isolates. The rows are the bacterial species (n next to the species name is the number of isolates tested); the columns are the 17 antibiotics tested. All 17 antibiotics are listed, including those to which no isolate was resistant. Abbreviations: CEF – cefoxitin, B-PEN – benzylpenicillin, OXA – oxacillin, GEN – gentamicin, CIP – ciprofloxacin, MOX – moxifloxacin, ERY – erythromycin, CLIN – clindamycin, LIN – linezolid, TEI – teicoplanin, VAN – vancomycin, TET – tetracycline, TIGE – tigecycline, FOM – fosfomycin, FUS – fusidic acid, RIF – rifampicin, TRIM – trimethoprim.

Regarding the prevalence of *Staphylococcus/Mammaliicoccus* bacteria in the samples, AMR and MDR prevalence in AST-tested isolates, and the summarised MAR index, the values and their comparisons between the two areas and the two host groups by Fisher’s exact tests are presented in Table 5 and Table 6, respectively. Prevalence data and MARi are visualised in Figure 5.

**Table 5.** Summarising the prevalence of *Staphylococcus/Mammaliicoccus* infected hosts (Bacterial prevalence), antimicrobial resistance (AMR), multidrug resistance (MDR), and Multiple Antimicrobial Resistance index (MARi) of the bacterial communities in the host groups (Cervidae and Suidae) of the two study sites (Somogy, SOM and Vas, VAS).

| Area | Host | Bacterial prevalence | AMR prevalence | MDR prevalence | MARi |
| --- | --- | --- | --- | --- | --- |
| SOM | Cervidae | 90.0% (27/30) | 50.0% (10/20) | 20.0% (4/20) | 6% |
| SOM | Suidae | 81.8% (18/22) | 56.25% (9/16) | 37.5% (6/16) | 9% |
| VAS | Cervidae | 40.0% (6/15) | 85.71% (6/7) | 14.29% (1/7) | 8% |
| VAS | Suidae | 66.67% (10/15) | 85.71% (12/14) | 21.43% (3/14) | 11% |

**Table 6.**
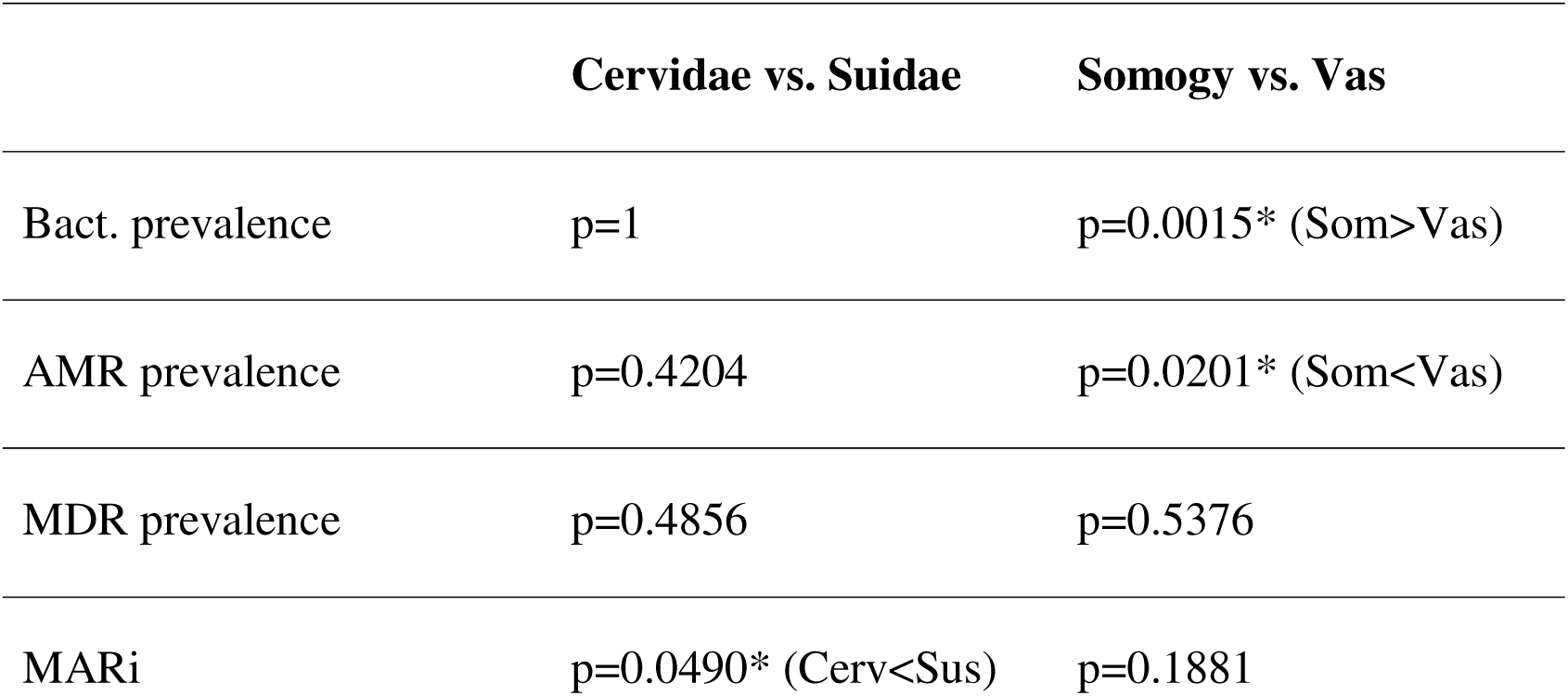
Statistical comparison of the prevalence values of *Staphylococcus/Mammaliicoccus* infected hosts (Bacterial prevalence), antimicrobial resistance (AMR), and multidrug resistance (MDR), and the multiple antimicrobial resistance index (MARi) of the bacterial communities between the host groups (Cervidae = Cerv and Suidae = Sus) of the two study sites (Somogy = Som and Vas = Vas). For comparison, Fisher’s exact test was applied. Asterisks mark significant differences between the compared bacterial communities. The directions of differences were marked with < and > signs.

**Figure 5.**
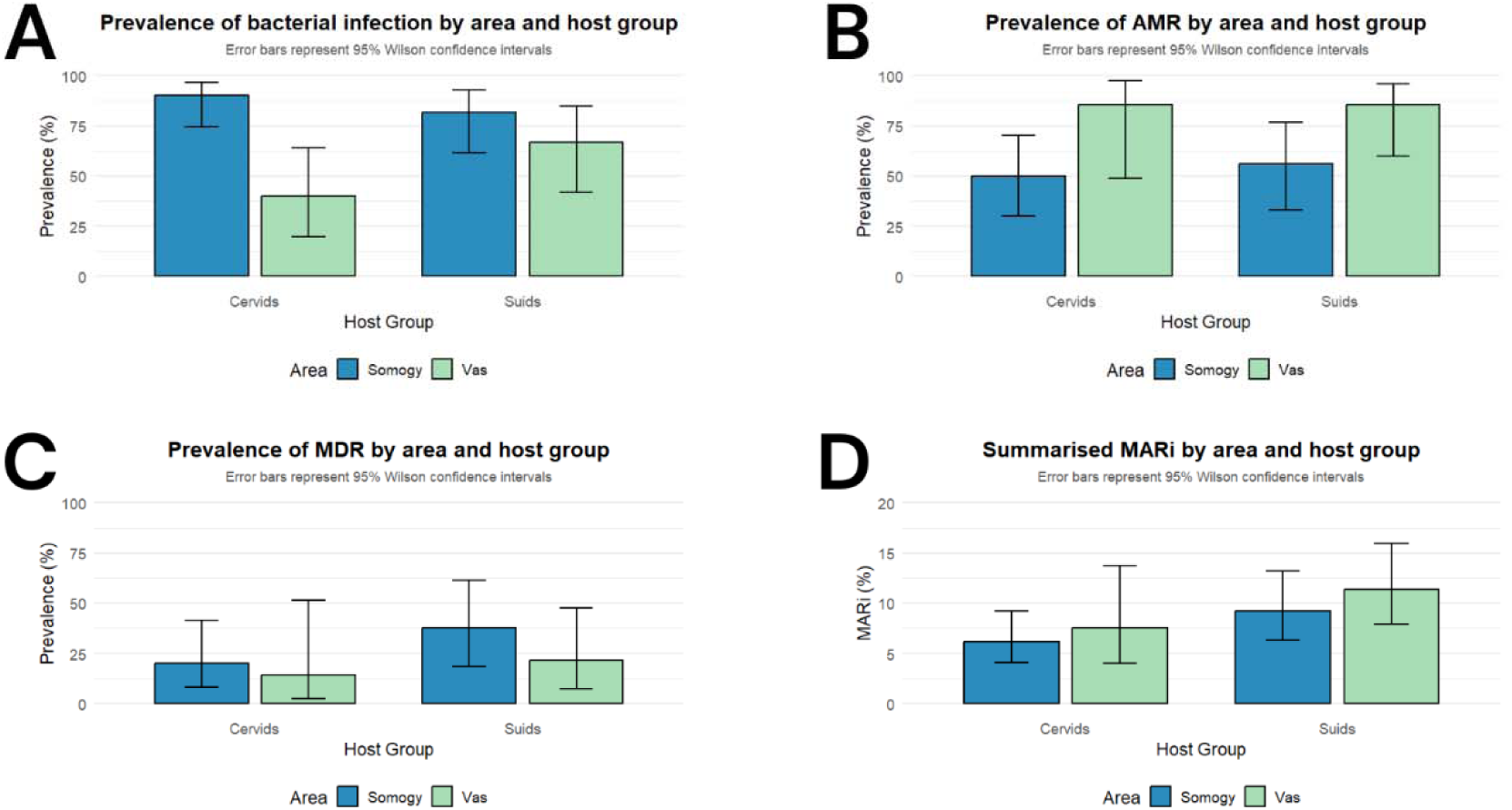
Visualisation of the comparison of the bacterial communities in the two host group populations (cervids and suids) of the two study sites (Somogy = SOM and Vas = VAS). A: prevalence of *Staphylococcus/Mammaliicoccus* bacterial infection (proportion of samples with an isolate of the genera *Staphylococcus* or *Mammaliicoccus*). B: prevalence of antimicrobial-resistant (AMR) isolates (isolates that are resistant to at least one antimicrobial). C: prevalence of multidrug-resistant (MDR) isolates (isolates that are resistant to at least three classes of antimicrobials). D: summarised Multiple Antimicrobial Resistance index (MARi) in the host population (proportion of resistant phenotypes among all phenotypes tested in antimicrobial susceptibility tests).

The Rényi profile of the landscape structure (Figure 6) yielded different pictures depending on the thematic resolution. At the level 1 (functional) resolution, both areas contained all six main categories (type richness is the same, α = 0: ln 6 = 1.79), so the difference clearly reflected uniformity: the profile of the Vas County area ran above the Somogy area on the entire scale (Shannon 1.14 vs 0.80; inverse Simpson 0.98 vs. 0.48; dominant weight [α = ∞] 0.80 vs. 0.26). Based on the dominance indicator, a single class, the forest, accounted for about 77% of the area in the Somogy landscape, and about 45% of the dominant arable land at the Vas site. The configuration indicators were in agreement: the landscape of the Vas site was more fragmented and richer in edges (ED 74.6 vs 48.5 m/ha) and patches (PD 24.1 vs 16.4 patches/100 ha).

**Figure 6.**
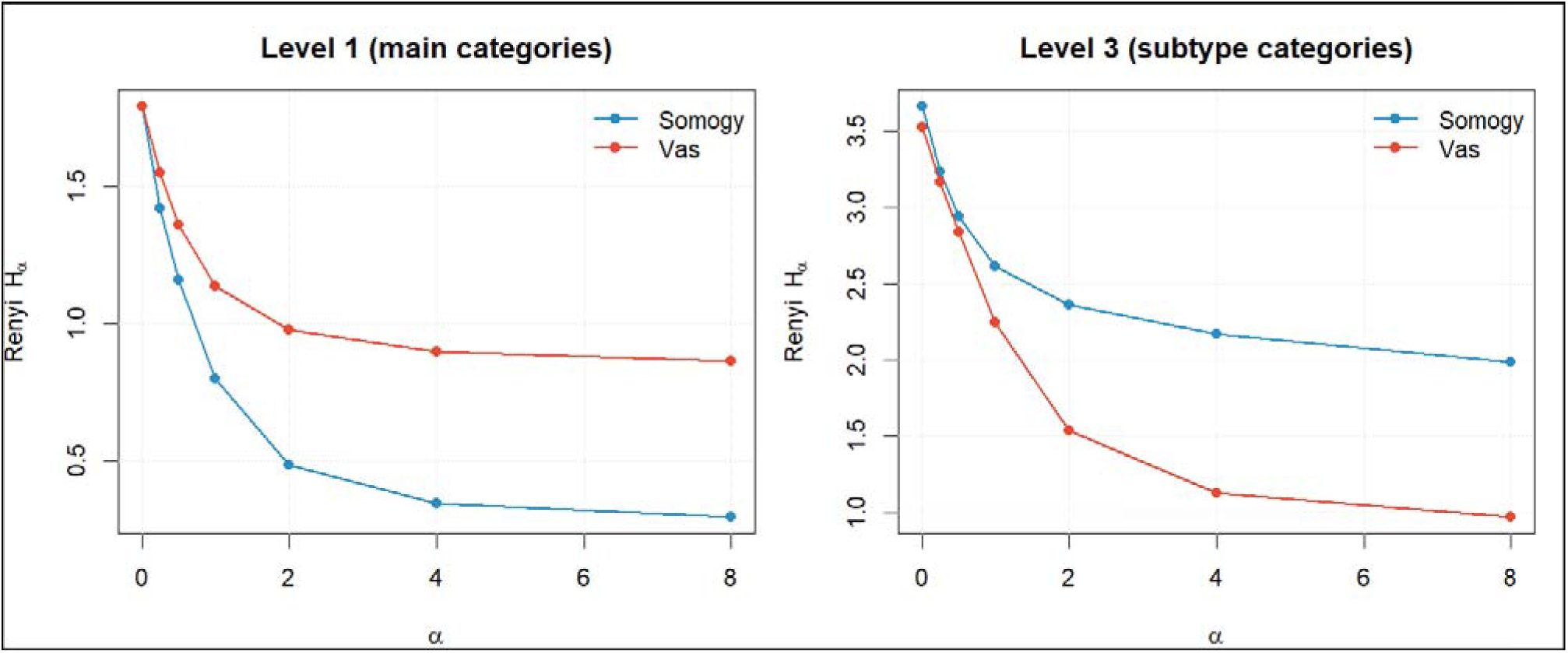
Rényi’s profile of the diversity of landscape structure (coverage) by area, at two thematic breakdowns (left: level 1, six main categories; right: level 3, subtypes). The “type” is the coverage category; its frequency is the territorial ratio; the x-axis is the α scale parameter, and the y-axis is the Rényi entropy. At level 1, the Vas site’s profile runs above the Somogy site’s profile on the entire scale (the landscape is more balanced; In Somogy, ∼77% forest dominance); at level 3, the order is reversed (due to the division of the Somogy forest block into subtypes), well illustrating that the ranking of landscape diversity depends on the thematic resolution.

At level 3 (subtype) resolution, the order of diversity was reversed: the Rényi profile of the Somogy landscape ran higher than the Vas landscape throughout the scale (Shannon 2.61 vs. 2.25; inverse Simpson 2.36 vs. 1.54; α = ∞: 1.75 vs. 0.85), while the PD (66.8 vs. 70.6 m/ha); and ED (153.3 vs. 161.5 patches/100 h) were practically equalised. The reason for the turnaround is that at this breakdown, the contiguous forest block of Somogy is divided into about a dozen native forest subtypes, making the landscape compositionally rich and seemingly fragmented, while a single class, the arable land, still dominates the area of Vas. This result highlights that a landscape diversity index, in itself, is not level-invariant: the same metric can even yield an opposite ranking, depending on the choice of thematic resolution. (Figure 6).

The habitat composition of the areas differed markedly. The landscape of Somogy was dominated by native forest (about 60% of the area), with a smaller proportion of agricultural land (∼10%), grassland (∼8.5%) and only ∼2.6% artificial surface; the landscape of Vas, on the other hand, was dominated by agriculture (arable land ∼45%, non-native plantations ∼16%, artificial surface ∼10.5%), and native forests appeared only in ∼12%. The average naturalness relative to area was 4.01 in the Somogy region and 2.45 in Vas (on a scale of 1–5). The internal composition of the forest stock confirmed this: about 78% of the forest in Somogy belonged to the type with native tree species, compared to the ∼40% proportion of non-native plantations in the forest of the Vas site (where the proportion of native types was only ∼31%).

Rényi entropy analysis of bacterial species composition in different areas and host groups showed that Somogy has higher bacterial diversity (species richness 14 vs 10; Shannon diversity 2.27 vs 2.00) than Vas. According to the host group, the Suidae profile was above Cervidae in all parameters (species richness 16 vs 12; Shannon 2.60 vs 2.09) (Figure 7).

**Figure 7.**
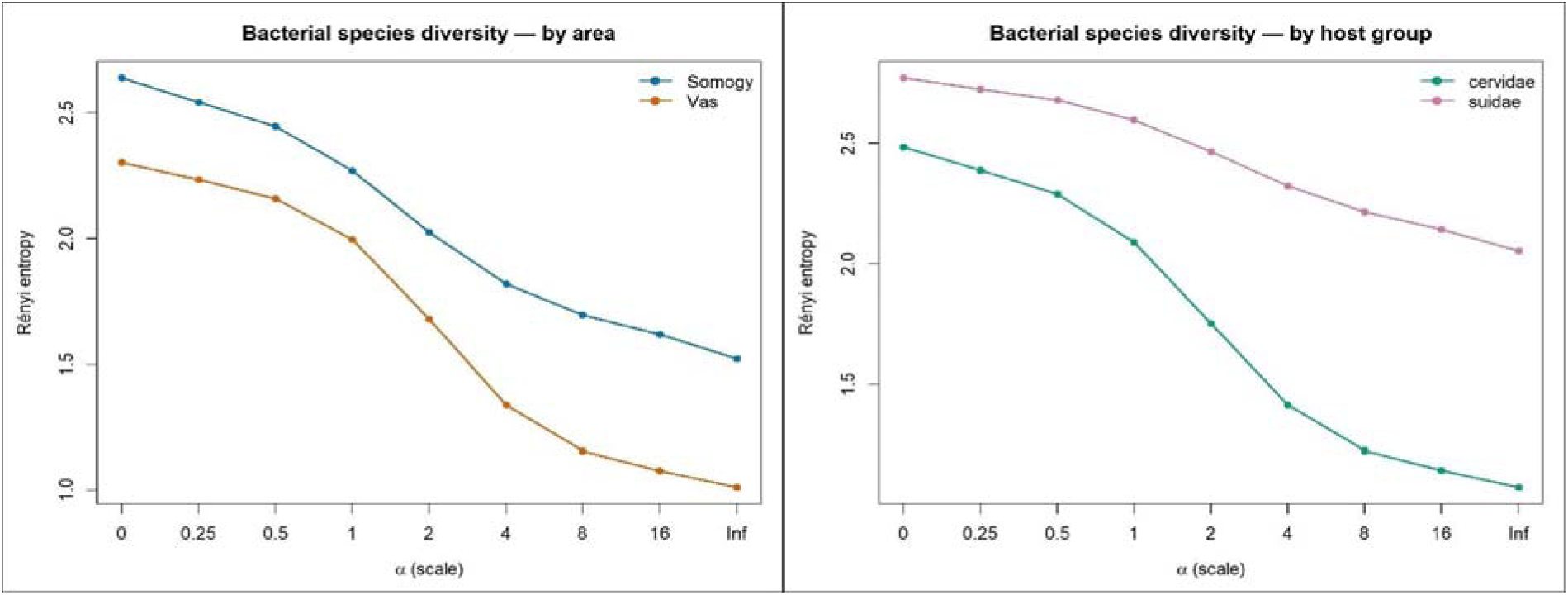
Rényi profiles of bacterial species diversity by area (left) and host group (right). The “type” is the bacterial species, and its frequency is the number of isolates. The x-axis is the α scale parameter (α = 0: species richness; α = 1: Shannon; α = 2: inverse Simpson; α = ∞: weight of the dominant species), and the y-axis is the Rényi entropy. By area, Somogy’s profile is above Vas on the entire scale (species richness 14 vs 10), and by host group, the Suidae profile is above Cervidae (16 vs 12). So, the diversity ranking is clear in both breakdowns (Somogy > Vas and Suidae > Cervidae).

The diversity of antibiotic resistance at the level of the active ingredient was almost the same by area: the two profiles crossed each other (the number of active substances showing resistance was 8 in Vas, 7 in Somogy; Shannon 1.65 vs 1.58), so the diversity of resistance between the active ingredients did not differ between the two areas (Figure 8). However, by host group, the Suidae profile was consistently above Cervidae (9 vs 7 active ingredients; Shannon 1.82 vs 1.50)

**Figure 8.**
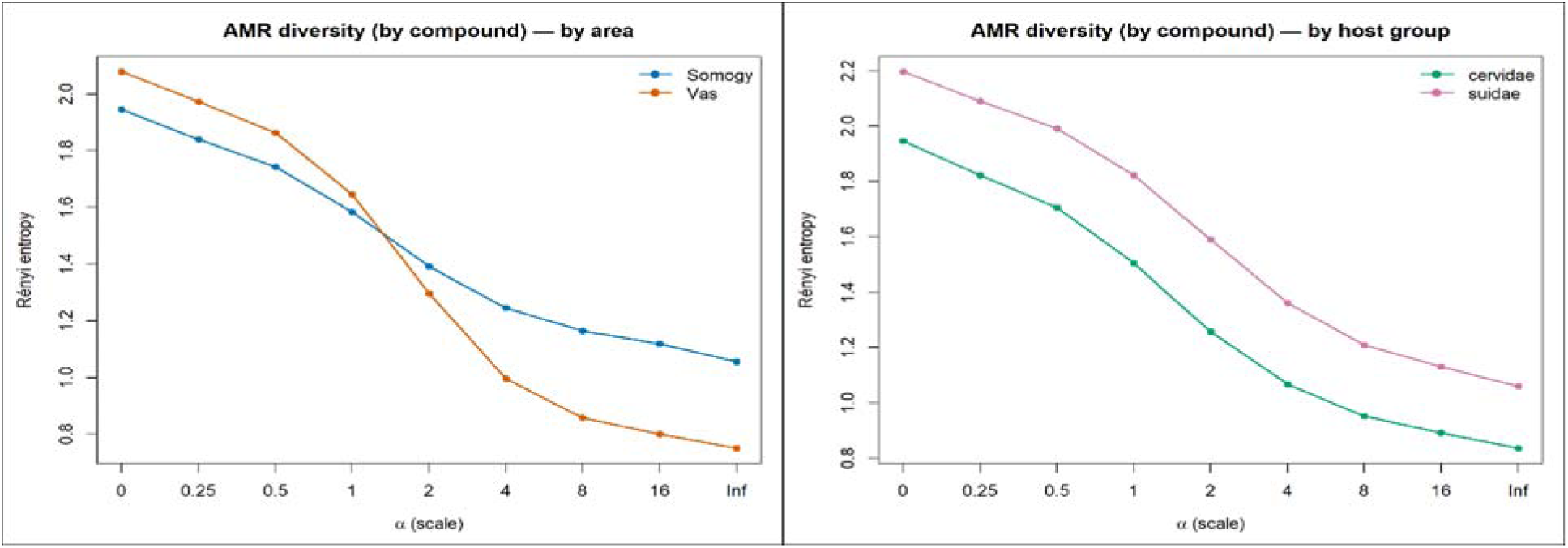
Rényi profiles of antibiotic resistance diversity by area (left) and host group (right).

The “type” is the antibiotic drug; its frequency is the number of resistant isolates. The x-axis is the α scale parameter (α = 0: drug richness; α = 1: Shannon; α = 2: inverse Simpson; α = ∞: weight of the dominant active ingredient); the y-axis is the Rényi entropy. By area, the two profiles intersect (between α ≈ 1–2), so the two areas are not comparable (resistance to more active substances appeared in Vas, but Somogy is higher in the dominant range). According to the host group, the profile of Suidae is above that of Cervidae across the entire scale, so the resistance diversity of Suidae is clearly greater.

## Discussion

### Isolation of Staphylococcus and Mammaliicoccus bacteria

This study aimed to test a potential surveillance method for estimating the AMR load of natural ecosystems. For this, lymph nodes were collected from carcasses of healthy wild ungulates harvested in the framework of regular hunting management. Utilising the halotolerant feature of *Staphylococcus* and *Mammaliicoccus* bacteria (Onyango & Alreshidi 2018), 10% salt was added to the pre-enrichment culture media. This step gained detection success between 40% and 90%. These findings align with previous studies on suids (Mann et al. 2015; Ravel et al. 2015) and ruminants (Belser et al. 2016; Hanlon et al. 2016) that revealed a remarkable prevalence of bacterial infections of lymph nodes along the digestive tract.

The ubiquitous distribution of *Staphylococcus* and *Mammaliicoccus* genera (Dhaouadi et al. 2026; Kochel-Karakulska et al. 2025), the presence in the lymphatic tissues of even visibly healthy animals (Mann et al. 2015; Linhares et al. 2015), and their relatively simple selective culturing (Rahman et al. 2026) make them good models to monitor AMR circulation between the three domains of health. Furthermore, European hunting technology enables large-scale sampling in the game meat processing plants, since wild boar carcasses are transported to the plants with heads, thereby including submandibular lymph nodes. On the other hand, our findings highlight the biological hazard of hunting offal remaining on the hunting ground. Viscera of both host groups and the heads of cervid carcasses are mostly dumped on-site, allowing scavengers and decomposing organisms to acquire potentially zoonotic bacteria and antibacterial resistance genes.

The lowest apparent prevalence (40%) of *Staphylococcus/Mammaliicoccus* bacteria was detected in the lymph nodes of red deer in the Vas area. Although behind this apparent value, the true prevalence can be substantially lower, it is high enough to conduct statistical analyses (e.g., comparison of bacterial communities).

### Landscape complexity and bacterial community

The Rényi profile analysis based on the main land cover categories of the two areas shows that the Vas site has higher landscape diversity. Although both have the same land cover types, the Somogy area is dominated by forest habitats (77% of the whole territory); therefore, Vas appeared to be a more balanced landscape mosaic. However, using sub-type categories (e.g., forest typology instead of “forest”), the analysis highlighted the finer structure of the landscape, resulting in an opposite Rényi profile. In parallel, the average naturalness, which expresses the degree of human influence in the two sites, demonstrates that the Somogy site is a more natural landscape (N=4.01) than the Vas site (N=2.45).

Landscape mosaics with plenty of small habitat patches are divided by an extended network of edges. These transitional habitats not only mix the species of the bordering habitats but also have their own edge-related species (Erdős et al. 2019). Therefore, the presence of edges might increase the effective plant species number, plant species richness, and the number of red-listed plant species (Pöpperl & Seidl 2021). This effect is supported by grazing, which is characteristic of the grasslands of the Vas site. Large herbivores at low density can support the ecological restoration of grasslands, contributing to the increase of both plant species richness and the prevalence of regionally rarer species (Bonavent et al. 2023). On the other hand, the Vas site is dominated by arable land, and a large proportion of its forest habitats are non-native robinia stands, resulting in lower average naturalness. The apparent landscape diversity is likely due to an artificial habitat fragmentation rather than richness in natural habitats. Furthermore, the permanent presence of domestic animals, beef cattle on the grasslands of the Vas site likely led to spill-over of both bacteria and AMR phenotypes of the domestic cycle. Therefore, summarising the analyses of all landscape metrics, the Somogy site was evaluated as a more diverse, more complex, and more natural landscape.

Comparing the Rényi profiles of the Somogy and the Vas bacterial communities, Somogy appeared to have a higher diversity that is also supported by Fisher’s exact test. The pair-wise comparison of the two communities revealed that the dissimilarity is predominantly due to *S. epidermidis*, the dominant bacterial species in the Vas area. Interestingly, this member of the *Staphylococcus* genus is highly adapted to humans. Although it is well-documented that *S. epidermidis* can be found in the microbiota of several animal and plant species (Chaudry & Patil 2016; Garcia-Gutierrez et al. 2020), the high occurrence at the Vas site, particularly in both host groups, suggests an anthropogenic influence.

On the other hand, mammaliicocci are characteristic of the Somogy site, with ten *M. sciuri*, two *M. vitulinus*, and one *M. lentus* isolates, while only one *M. vitulinus* isolate was detected at the Vas site. The genus *Mammaliicoccus* was separated from the genus *Staphylococcus* based on a phylogenomic analysis using 16 rRNA sequencing. As a result, *Staphylococcus sciuri*, *S. vitulinus*, *S. lentus*, *S. fleuretti*, and *S. stepanovici* was reassigned to the novel genus *Mammaliicoccus* (Madhaiyan et al. 2020). *Mammaliicoccus sciuri*, the type species of the genus, is regarded as a basal species in the Staphylococcaceae family (de Moura et al. 2023). Mammaliicocci are also considered a natural reservoir of antimicrobial resistance. Besides the *mecA* (responsible for methicillin resistance), mammaliicocci possess several other resistance genes that enable them to survive the effects of, e.g., fusidic acid, clindamycin, and ciprofloxacin (Dhaouadi et al. 2026). A study that investigated the temperate phages in *M. sciuri* revealed an unexpectedly high phage diversity, suggesting a long-standing co-evolutionary process between the bacterial species and its parasite (Cherbuin et al. 2025). In addition, the genomic profiling of *M. sciuri* demonstrated that its CRISPR-Cas system (responsible for the defence against mobile genetic elements) constitutes a small part of the genome, particularly compared to the number of resistance genes, contributing to the adaptability of mammaliicocci (de Carvalho et al. 2024). These features promote survival across a wide range of matrices worldwide (Dhaouadi et al. 2026).

Research data suggest that the genus *Mammaliicoccus* occupies a basal position within the family Staphylococcaceae, closest to the soil-inhabiting ancestors of the family. Although the only *Mammaliicoccus* isolate of the Vas site was found in a wild boar sample (explainable by soil-associated feeding habits of the species), at the Somogy site, mammaliicocci were found in both host groups equally. The *Mammaliicoccus* diversity of the Somogy site is relatively high, since three of the five currently known species of the genus were detected here. Therefore, future research based on a larger sample size is needed to highlight the interestingly frequent occurrence of mammaliicocci in the Somogy area.

The second most dominant taxon in the Somogy area was *S. hyicus*, the causative agent of exudative epidermitis in pigs (Park et al. 2013). Besides domestic pigs, this coagulase positive bacterium occurs also in healthy wild boars (Romero-Salmoral et al. 2025). In our study, *S. hyicus* was found in both host groups with approximately the same frequency, consistent with the findings of a Central Italian study where this species was detected in the nasal specimens of fallow deer (Cagnoli et al. 2024). Interestingly, this species was completely absent from the Vas site, contrary to the presence of wild boars. This finding is similar to the experiences in Spain, where the prevalence of *S. hyicus* carriage in wild boars was also low, 2.7% (Mama et al. 2019a).

Besides bacterial diversity, in the Somogy ungulate populations, more bacterial isolates were obtained with the same method compared to Vas. Together with the high frequency of mammaliicocci, this phenomenon is also worth investigating. Since the most conspicuous difference between the two sites is the predominance of forest habitats and higher naturalness at the Somogy site, future research would focus on silvatic habitat as a potential reservoir of the genus *Mammaliicoccus* and a source of bacterial diversity.

### Landscape complexity and AMR pattern

Rényi diversity profile analysis showed that the AMR phenotype richness was higher at the Vas site, while Somogy had more isolates with the generally dominant phenotypes, such as B-PEN, FUS, and CLIN. The statistical comparison revealed that the Somogy site has a more heterogeneous AMR profile, which means that the distribution of resistant phenotypes among the isolates is highly variable. Two phenotypes, TET and ERY, are completely absent from Somogy, while two phenotypes, CLIN and FOM, occur less frequently in the Somogy area than in Vas. Furthermore, the prevalence of AMR isolates, possessing resistance to at least one antimicrobial, was significantly higher in Vas than in Somogy. These findings suggest that the antimicrobial pressure at the Vas site is higher compared to Somogy.

This hypothesis is also supported by the fact that CLIN-and FUS-resistant phenotypes in Somogy are exclusively associated with *M. sciuri*, a bacterial species with well-documented intrinsic resistance to CLIN and FUS. As a natural reservoir of several resistance genes, *M. sciuri* can show phenotypic AMR without anthropogenic antimicrobial pressure (Dhaouadi et al. 2022).

The importance of B-PEN and TET resistant phenotypes was high at the Vas site. These two antimicrobials have been the most important veterinary drugs for decades (van Rennings et al. 2015). The presence of the resistant phenotypes indicates anthropogenic impact. Interestingly, half of the B-PEN phenotypes in Somogy were detected in methicillin-susceptible *M. sciuri* isolates, contrary to the fact that this species is considered the origin of the *mecA* (methicillin resistance coding) gene of the Staphylococcaceae family. In the presence and expression of the *mecA* gene, bacteria show resistance to most beta-lactams, including B-PEN, OXA, and FOX (Schwendener & Perreten 2022). Our finding suggests that *M. sciuri* isolates of Somogy might have lost their *mecA* genes or their genes were “silent”. However, the B-PEN-resistant phenotypes of *M. sciuri* suggest an alternative genetic background for B-PEN resistance, e.g., the *blaZ* gene (Schwendener & Perreten, 2022), which needs molecular genetic investigations to clarify. Apart from these B-PEN-resistant *M. sciuri* isolates, the other seven B-PEN phenotypes of Somogy were distributed between *S. aureus* (N=2), *S. xylosus* (N=2), *S. equorum* (N=1), *S. gallinarum* (N=1), and *S. saprophyticus* (N=1), while 22 isolates showed susceptibility to B-PEN. In contrast, the Vas-originated isolates showed an 81% prevalence of the B-PEN phenotype, indicating a higher selection pressure by the antimicrobial.

In addition to the traces of veterinary impact, the exclusive presence of the ERY phenotype at the Vas site suggests an AMR load of public health origin. A previous study found that the increasing use of macrolide, lincosamide, and streptogramin B (MLSB) antibiotics in human patients as alternatives to beta-lactams correlates with the emergence of ERY resistance (El Mammery et al. 2023). Together with the finding that the Vas site is dominated by *S. epidermidis*, a human-adapted bacterial species, the detection of the ERY phenotype strengthens the hypothesis that the Vas site is under a stronger anthropogenic impact than the other study site.

Interestingly, the *S. hyicus* isolates (N=9), which were found only at the Somogy site in both host groups, were susceptible to all tested antibiotics. This was consistent with previous findings in wild boars in Spain (Mama et al. 2019a) but contrasted with the findings in fallow deer in Italy, where a 41.2% B-PEN resistance rate was detected in this bacterial species (Cagnoli et al. 2024).

Regarding MDR isolates, the Somogy site showed a higher prevalence, although this difference was not statistically significant. This phenomenon can be attributed to the high number of *M. sciuri* isolates, which likely possess intrinsic resistance features. This finding aligns with the fact that the summarised MARi of the Vas site was higher, suggesting a more constant circulation of AMR in the Vas area. In contrast, approximately half of the AMR phenotypes at the Somogy site are likely due to the intrinsic resistance associated with *M. sciuri*.

Methicillin resistance, indicated by the OXA and FOX resistant phenotypes, was detected at both sites (Somogy: N=2; Vas: N=1). The statistical comparison showed that this difference was non-significant. This feature was detected in *S. xylosus* and *S. equorum* in Somogy, and *S. arlettae* in Vas. None of the *S. aureus* isolates showed MRSA phenotypes in contrast to previous experiences (Monecke et al. 2016; Ramos et al. 2022). Among *S. aureus* isolates, only one MDR phenotype (resistant to B-PEN, CLIN, and TET) was detected at the Vas site. In comparison, *S. aureus* of the Somogy site showed resistance to B-PEN (N=2) and FUS (N=2).This narrow resistance profile of wildlife-originated *S. aureus* is uncommon compared to other European research that found various resistance genes in wild ungulates (Monecke et al. 2016).

### Bacterial diversity and AMR pattern in Suidae and Cervidae

Comparing the bacterial communities associated with the two host species showed that Suidae carry a more diverse bacterial community than Cervidae. Interestingly, the prevalence of identifiable *Staphylococcus/Mammaliicoccus* isolates was higher in Cervidae; although this difference was not significant. Considering the feeding ecology of wild boars, like rooting and scavenging (Drimaj et al. 2025), it was expected that suid hosts accumulate bacteria with high prevalence and diversity. Our finding on high *Staphylococcus/Mammaliicoccus* diversity is in parallel with a study conducted in wild boars in Spain (Mama et al. 2019b). Interestingly, the detectable prevalence of *Staphylococcus/Mammaliicoccus* bacteria at the Somogy site was higher in Cervidae than in wild boars. This phenomenon is likely due to our selective culture technique focused on halotolerant bacteria, thereby overlooking concurrent bacterial taxa that could be highly diverse owing to the soil-associated feeding behaviour of suids (Drimaj et al. 2025; Belardi et al. 2026).

Statistical comparison of the two host groups determined only one bacterial species, *S. aureus*, which was characteristic of Cervidae. This finding aligns with other studies that detected a high prevalence of *S. aureus* (Luzzago et al. 2022; Ramos et al. 2022) and the occurrence of MRSA (Monecke et al. 2016) in red deer. Considering that cervids are herbivores, being at a lower level of trophic webs, they have a lower chance of accumulating pollutants than the omnivorous wild boars (Chen & Li, 2026). A study conducted in the Central Italian Alps hypothesises that increasing deer density and their occurrence in anthropogenic landscapes led to the accumulation of human-adapted bacteria (Luzzago et al. 2022). However, the researchers did not find methicillin-resistant *mecA* and *mecC* genotypes, similar to our experience on the complete absence of FOX and OXA phenotypes, which challenges the theory of anthropogenic origin. Furthermore, the high predominance of deer-derived *S. aureus* isolates suggests that this bacterial species has strains adapted to Cervidae; although this hypothesis needs a comprehensive molecular investigation to elucidate.

Surprisingly, the theoretically swine-adapted *S. hyicus* was isolated from both cervid and suid hosts in equal proportion; however, only within the Somogy study area. Besides *S. aureus*, this species belongs to the coagulase-positive staphylococci. These bacteria are presumed to be pathogenic due to their ability to coagulate the extracellular fluid in the host’s tissues to avoid the immune system (González-Martín et al. 2020). However, in our study, bacteria were isolated from visibly intact lymph nodes, which contradicts the pathogenicity of the detected *S. hyicus* isolates. In addition, all these isolates were susceptible to all tested antimicrobials; therefore, we hypothesised that the *S. hyicus* population of the Somogy site could be a natural commensal with low invasive ability. In the future, a comparison to swine farm strains based on molecular techniques would highlight the role of these bacteria within the currently investigated hosts and their habitat.

Regarding AMR phenotypes, all indicators, such as prevalence of AMR and MDR, and MARi, showed higher values in Suidae than in Cervidae, with an interesting anomaly at the Vas site where the prevalence of AMR (isolates with at least one AMR phenotype) was equal in the two host groups. Although only the summarised MARi was significantly higher in Suidae than in Cervidae, wild boars appear to accumulate more resistant phenotypes. This would be explainable by the omnivorous and scavenging lifestyle of wild boars (Chen & Li 2026; Drimaj et al. 2025). However, it is worth noting that conspicuous AMR accumulation by wild boars was documented in association with other bacterial taxa, like *Escherichia coli* (Höcketstaller et al. 2025), while studies dealing with staphylococci demonstrated that Cervidae appear to be more important reservoirs of these bacteria and their resistant genotypes than wild boars (Cagnoli et al. 2024; Mateus-Vargas et al. 2022; Plaza-Rodríguez et al. 2021; Ramos et al. 2022).

Regarding host groups, our main findings suggest that deer species are reservoirs of *S. aureus*, while wild boars accumulate more AMR genotypes than cervids. However, these hypotheses need to be supported by a larger sample size and molecular genetics in future research.

### Landscape ecology and the epidemiology of AMR

This One Health study combined landscape ecology and bacteriology to elucidate some potential drivers of AMR circulation in wild ungulates. Landscape characteristics, such as diversity, patch density, edge density, and naturalness, were applied to compare the two study sites. Based on fine-scale landscape categories, it was demonstrated that one of the study sites is under a stronger anthropogenic impact than the other.

Comparing the bacterial communities and the AMR phenotypes of the two sites, some interesting properties were detected. Regarding the more human-influenced environment, the predominance of *S. epidermidis*, a human-adapted species, characterised the local *Staphylococcus/Mammaliicoccus* community. Furthermore, this community showed a diverse AMR pattern with the frequent occurrence of B-PEN and TET phenotypes. On the other hand, the bacterial community of the more natural landscape was dominated by *M. sciuri*, and most of the AMR phenotypes were associated with this bacterial species. Since *M. sciuri* possesses several intrinsic resistance genes, it is likely that at least a part of the local resistance is independent of anthropogenic influence. Another interesting finding was the high prevalence of pan-susceptible *S. hyicus*, a coagulase-positive species, equally in both host groups within the more natural landscape, while it was completely absent from the anthropogenic environment.

The findings of this study suggest that the *Staphylococcus* and *Mammaliicoccus* genera would be effective models to evaluate the human impact on bacterial communities. Besides their ubiquitous distribution and simple selective culture, the composition of their communities seems to reflect the level of hemeroby. Although a study limited to two study sites is not sufficient for drawing definitive conclusions, future research would investigate the potential of the *Mammaliicoccus* genus as an indicator of naturalness. Similarly, a comparative molecular genetic study would explore the differences between *S. hyicus* isolates from pig farms and natural environments, since this species was also detected only in the landscape with lower human impact.

Apart from *S. aureus*, which appeared to be associated with deer species, no other relevant difference was determined between the two host groups, enabling both to be potential sentinels to detect *Staphylococcus/Mammaliicoccus* bacteria. Regarding AMR patterns of the two hosts, the significantly higher MARi in wild boars was the only attribute, which suggests that Suidae would be a better target taxon for future AMR monitoring.

This study has several limitations. Data were collected from only two study sites; therefore, the association between landscape features and bacterial communities cannot be ascertained by statistical analyses. In a future study, an appropriate number of study sites would provide a dataset suitable for statistical modelling. In the absence of molecular genetic investigations, the backgrounds of AMR phenotypes could not be ascertained. Furthermore, the role of the *Mammaliicoccus* genus and *S. hyicus* in natural ecosystems would need a comprehensive genetic investigation. Based on the current study, the hemeroby indicator role of these taxa is just an interesting hypothesis, which needs further research to prove.

## Conclusion

This study aimed to analyse the association between landscape complexity and the local AMR load and evaluate the potential human impact in the background. The anthropogenic influence on landscapes was determined by calculating landscape diversity, patch density, edge density, and naturalness. Using *Staphylococcus/Mammaliicoccus* bacteria as models, the composition and AMR profile of the bacterial communities of a more anthropogenic and a more natural landscape were analysed. The results suggest that the predominance of the *Mammaliicoccus* genus and pan-susceptible *S. hyicus*, and a heterogeneous AMR pattern might be indicators of naturalness. On the other hand, the predominance of human-associated *S. epidermidis* and high richness of AMR phenotypes might indicate stronger human impact.

